# MechanoProDB: A Web Based Database for Exploring the Mechanical Properties of Proteins

**DOI:** 10.1101/2023.11.30.569409

**Authors:** Ismahene Mesbah, Bianca Habermann, Felix Rico

**Affiliations:** Aix Marseille Univ, CNRS, INSERM, LAI, Turing center of living systems (CENTURI), Marseille, France; Aix Marseille Univ, CNRS, IBDM UMR7288, Turing Center of Living Systems (CENTURI), Marseille, France

## Abstract

The mechanical stability of proteins is crucial for biological processes. To understand the mechanical functions of proteins, it is important to know the protein structure and mechanical properties. Protein mechanics is usually investigated through force spectroscopy experiments and simulations that probe the forces required to unfold the protein of interest. While there is a wealth of data in the literature on force spectroscopy experiments and steered molecular dynamics simulations of forced protein unfolding, this information is spread and difficult to access by non-experts. Here we introduce MechanoProDB, a novel web-based database resource for collecting and mining data obtained from experimental and computational works. MechanoProDB provides a curated repository for a wide range of proteins, including muscle proteins, adhesion molecules and membrane proteins. The database incorporates relevant parameters that provide insights into the mechanical stability of proteins and their conformational stability such as the unfolding forces, energy landscape parameters and contour lengths of unfolding steps. Additionally, it provides intuitive annotations of the unfolding pathways of each protein, allowing users to explore the individual steps during mechanical unfolding. The user-friendly interface of MechanoProBD allows researchers to efficiently navigate, search and download data pertaining to specific protein folds or experimental conditions. Users can visualize protein structures using interactive tools integrated within the database, such as Mol^*^, and plot available data through integrated plotting tools. To ensure data quality and reliability, we have carefully manually verified and curated the data currently available on MechanoProDB. Furthermore, the database also features an interface that enables users to contribute new data and annotations, promoting community-driven comprehensiveness. The freely available MechanoProDB aims to streamline and accelerate research in the field of mechanobiology and biophysics by offering a unique platform for data sharing and analysis. MechanoProDB is freely available at https://mechanoprodb.ibdm.univ-amu.fr.

## INTRODUCTION

Mechanical forces are involved in almost all biological processes and play a significant role, for example during muscle contraction^1^, hearing ^2^, cell adhesion^3^, protein folding at the ribosomal exit tunnel^4^ and molecular motor activity^5^. Proteins are the main participants in these events. These nano-elements support, transmit or exert mechanical force. Therefore, studying the mechanical properties of proteins helps us understand their structure-function relationship. The resistance to unfolding in response to a mechanical force, also called mechanical stability, is a physiologically important parameter that permits the molecule to remain folded under mechanical stress. The mechanical stability of single proteins is commonly probed by forced unfolding experiments ^6,7^. Methodologically, techniques like atomic force microscopy (AFM)^8^, magnetic and optical tweezers^9^ allow us to probe unfolding forces at the single-molecule level through single-molecule force spectroscopy (SMFS) experiments. In SMFS unfolding experiments, single proteins are grabbed from their end using the probe and pulled at a constant speed. This applies an increasing force to the protein until it eventually unfolds at a given force. Protein unfolding is a stochastic process that depends on the applied pulling or loading rate, i.e. the rate at which the force is applied (pN/s). Therefore, average rupture forces are generally reported for different loading rates in the so-called dynamic force spectrum (DFS, force versus loading rate). Fitting theoretical models to the DFS allows us to extract energy landscape parameters such as the distance to the transition state *x*_β_ and intrinsic unfolding rate *k*_*off*_ ^10^. In addition, computational techniques such as steered molecular dynamics simulations (SMD)^11^ enable atomic level understanding of these molecular processes by mimicking SMFS experiments. Both experimental and computational approaches have been applied extensively to probe the mechanical stability of proteins in recent years, reporting unfolding forces at different pulling velocities or loading rates on hundreds of proteins^7^. However, accessing and understanding the vast amount of available experimental and computational data of protein unfolding remains challenging for non-experts.

While there are excellent reviews compiling a considerable amount of experimental forced unfolding data on different protein systems that inspired our work, compiled data is not available in the form of an accessible database^7,12^. The only existing database in the field is the Biomolecule Stretch DataBase (BSDB)^13^. BSDB mainly stores synthetic data obtained from a Go-like model^14^. Experimental data extracted from a literature survey is also available in BSDB^13^, however the compiled parameters are limited to protein length, unfolding forces, velocities and CATH references^15^ and to works published between 1997 and 2009. However, as unfolding forces depend on the experimental conditions, information about the energy landscape and kinetic parameters is missing. Moreover, an interactive and more user-friendly interface would allow the user to, for example, filter the data using keywords, broadening its usage.

To fill this gap and to allow users to search, filter, interact and contribute with data, we introduce MechanoProDB, a web-based database that will serve as a comprehensive and curated resource for experimental and computational forced unfolding of proteins. MechanoProDB was conceived and developed in response to an increasing demand within the molecular mechanobiology and single molecule biophysics communities to have a centralized, freely accessible, and updated resource.

MechanoProDB gathers data on different proteins, including muscle proteins, adhesion molecules, and membrane proteins that have been probed mechanically. MechanoProDB holds information about the mechanical stability of proteins, such as unfolding forces, velocities, loading rates, free energy landscape parameters, and contour lengths of unfolding steps. Other relevant information such as protein sequence, protein 3D structure, functional class and SCOP/SCOPe^16,17^ structural fold is also provided within MechanoProDB. Furthermore, we introduced intuitive manual annotations of the unfolding pathway that facilitate retrieving data using regular expressions.

The user-friendly interface enables researchers to efficiently explore, search, and download data related to specific protein folds or experimental conditions. Moreover, interactive tools and plugins allow users to visualize protein structures, while integrated plotting tools allow them to easily represent data. To ensure the reliability and quality of the database, the available data has undergone meticulous manual verification and curation. Our manual annotation of the unfolding process, unfolding pathways, and structural features enhance the depth and accuracy of available high-quality experimental and computational information, which is not available in other resources.

## MATERIAL AND METHODS

### DataBase structure, features and content

The MechanoProDB database is composed of one main table: MPDB_Proteins which contains the collected and annotated data from literature. Contributions from users are manually annotated and inserted within MPDB_Proteins. MechanoProDB was developed in HTML5/PHP 8.1 and MySQL 8.0. The main JavaScript^18^ DataTable of the database contains 34 columns storing both mechanical and structural information. Visualization of protein structure and sequence simultaneously is incorporated using the Mol*^19^ plugin. Complementary information about protein structure and function is extracted from the protein data bank (PDB) using an automated python script. This information includes SCOP structural annotation, functional classification, organism, and protein length.

Additionally, statistics of the database content is represented using the Plotly plugin of javascript which enables the user to interact, show/hide and even change the way data is plotted.

### Contributions from users

In MechanoProDB, users are given the possibility to contribute data and provide feedback. A dedicated page was built for this purpose. To facilitate this process, users who want to submit new results of proteins having a solved experimental structure, will need to provide only the Protein DataBank Identifier (PDB ID), as structural information, in addition to the unfolding forces with their respective velocities or loading rates. Other complementary information about structure and function such as SCOP annotation, functional classification, organism, and protein length, will then be extracted from the Protein DataBank (PDB) using our automated Python script. The contribution page takes as input measurements of the unfolding forces with their corresponding velocities and/or loading rates. For larger datasets, a template is available to upload data as a csv or any tabulated file containing data points.

## RESULTS

### DataBase structure, features and content

MechanoProDB is publicly available and freely-accessible at https://mechanoprodb.ibdm.univ-amu.fr. At the time of writing this paper, a total of 82 manually annotated entries are saved within the main datatable of MechanoProDB. Each entry stores mechanical parameters about a specific protein extracted from the literature and obtained under specific experimental or simulated conditions such as pulling velocity and/or loading rate, temperature, pulling geometry or even potential hydrogen (pH) condition. Each protein entry is displayed in one row that has 30 columns storing a variety of relevant information ranging from structural and functional to mechanical (**Figure 1**). One important aspect of MechanoProDB is that it includes SCOP^16^ annotations for the stored proteins. SCOP is a well-known resource that provides evolutionary information and structural classification of proteins which is useful for clustering and analysis of structural data and is manually annotated. Importantly, within MechanoProDB, these structural annotations are linked to the SCOPe database^17^ which is the extended version of SCOP that contains new protein structures. This link allows users to access a dedicated web-page in the SCOPe database, which provides detailed information about the structural fold of the protein as well as UniProt information. This link is valuable for users who want to further investigate the structural, functional and even evolutionary characteristics of the proteins of MechanoProDB. Furthermore, in our database, we provide mechanical clamps described previously^20–22^. These indicate which parts of the protein are bearing the force. A link to the web-based topology diagram^23^ is also provided when clicking at each mechanical clamp.

**Figure 1:**
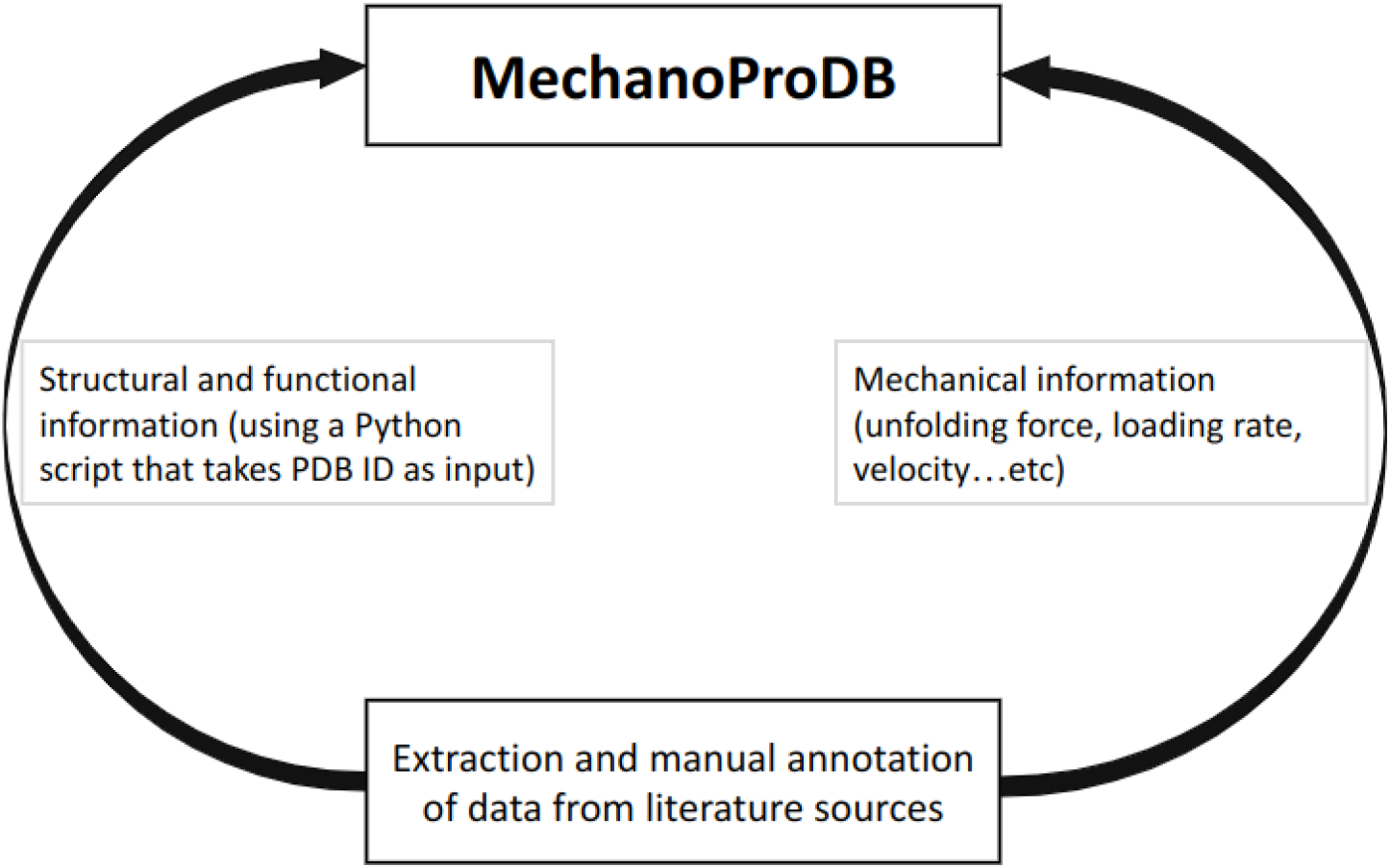
Schematic representation of MechanoProDB. As a first step, data is extracted and annotated from literature sources. The structural and functional information is retrieved from the PDB using a Python script when the PDB accession code is available or introduced manually by the user.

Detailed information about the original publication reporting the mechanical information is also provided in MechanoProDB. For example, internal links show relevant figures of force-distance curves and dynamic force spectra (force versus pulling speeds or loading rates). Moreover, if the PDB accession code of the protein or a close ortholog is available, MechanoProDB provides external links for each protein to its Protein DataBank (PDB) page in addition to the bibliographic reference from which mechanical data have been extracted and annotated. All the information of the database can be exported in a csv tabulated file to enable users to store and analyze data locally.

Data is stored within a DataTable variable which enables users to filter, search and manipulate data subsets. Through the intuitive “collections” button (**Figure 2)**, users can show/hide information of their interest, which enhances the efficiency of data visualization, extraction and analysis.

**Figure 2:**
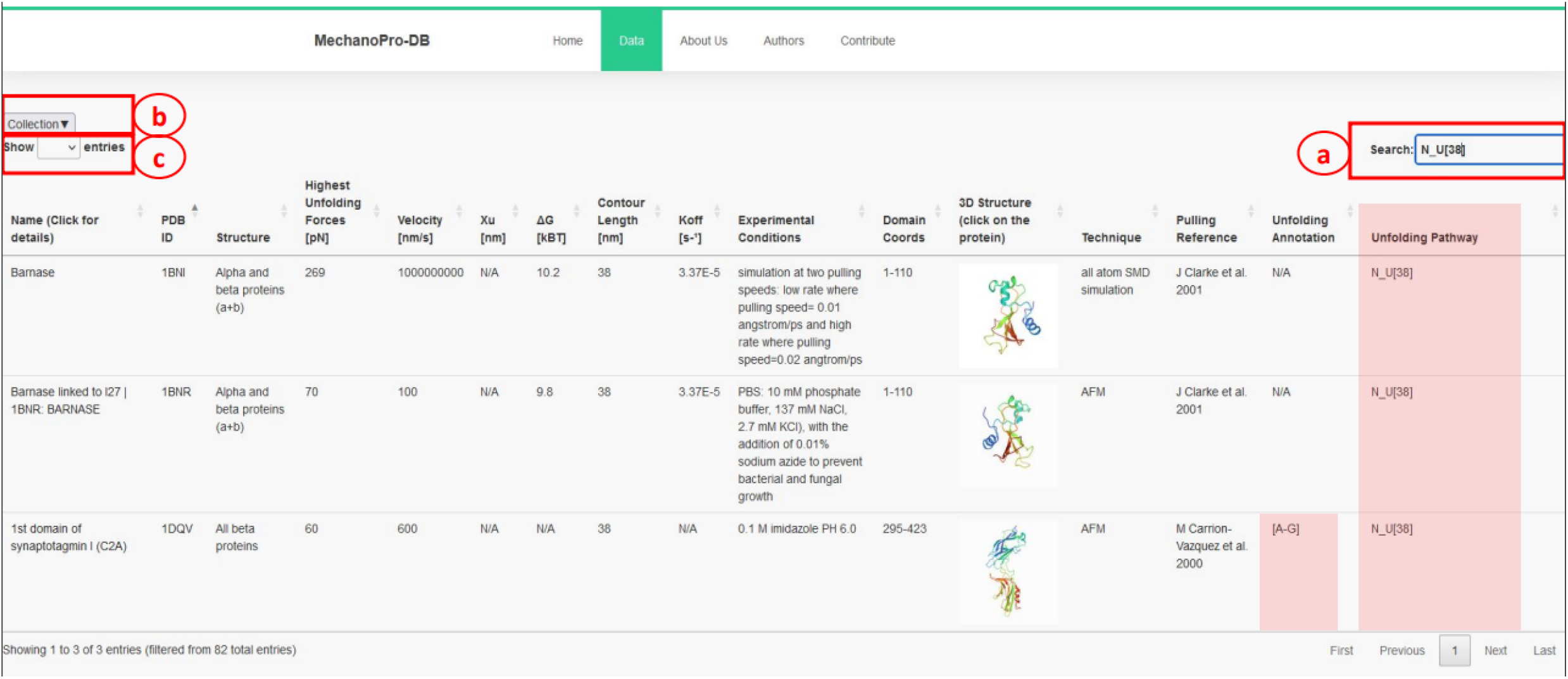
Screenshot of MechanoProDB’s main page. An example of searching for proteins with single step unfolding and a contour length of 38 nm is shown in (a). The unfolding pathway of interest here: *N* − *U*[38] is similar for three proteins and the available unfolding annotation of C2A is highlighted in red. The main table displays information including the PDB ID, unfolding forces, contour lengths, experimental conditions, domain amino acid coordinates, and unfolding process annotations. (b) is the collection button that enables users to show/hide parameters of interest and also allows downloading the stored data to their local machine. (c) shows the button enabling users to show the maximum number of entries displayed.

### Mechanical information and annotations

A unique feature of MechanoProDB is the manual annotation provided to describe the unfolding process and pathway for each protein entry. The unfolding pathway indicates the different steps before the unfolded state is reached with the contour length of each step. This annotation is written in a regular expression manner in which we report the unfolding step with its corresponding contour length. The regular expression pattern always starts with the native state (*N*) and ends with the unfolded state (*U*), some proteins require more than one step to unfold and therefore have intermediate steps (*I*).

The annotation of the unfolding pathway of a given protein respects the following structure:

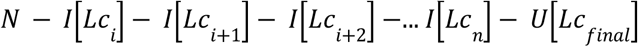

where, *Lc*_*i*_ is the contour length of the unfolded polypeptide chain after intermediate state (*I*) and *Lc*_*final*_ is the maximal physical extension of the chain after complete unfolding.

This regular expression process facilitates retrieving data for further analysis. Consequently, a user can, for example, extract all the proteins that unfold in one step (from native to unfolded state) with a contour length of 38 nm by simply writing *N*_*U*_[38]within the search entry (**Figure 2**).

In addition to the unfolding pathway annotation, we provide, when available in the original reference, which parts of the protein are involved in the unfolding process. **Figure 2** shows an example of annotation of the unfolding process of the first domain of synaptotagmin I (C2A)^24^. The annotation reported as [A-G] suggests that β-strands A and G are pulled apart during unfolding and are the main responsible substructures involved in supporting resistance to force. Another example is given in **Figure 3** with the I27 (previously named I91) module of titin. This domain has been studied extensively, and several entries obtained from different references are saved within MechanoProDB. The annotation provided for the unfolding process of I27 is [A-B] [A’-G] which indicates that the protein is unfolded once β-strands A and B break apart followed by β-strands A’ and G.

**Figure 3:**
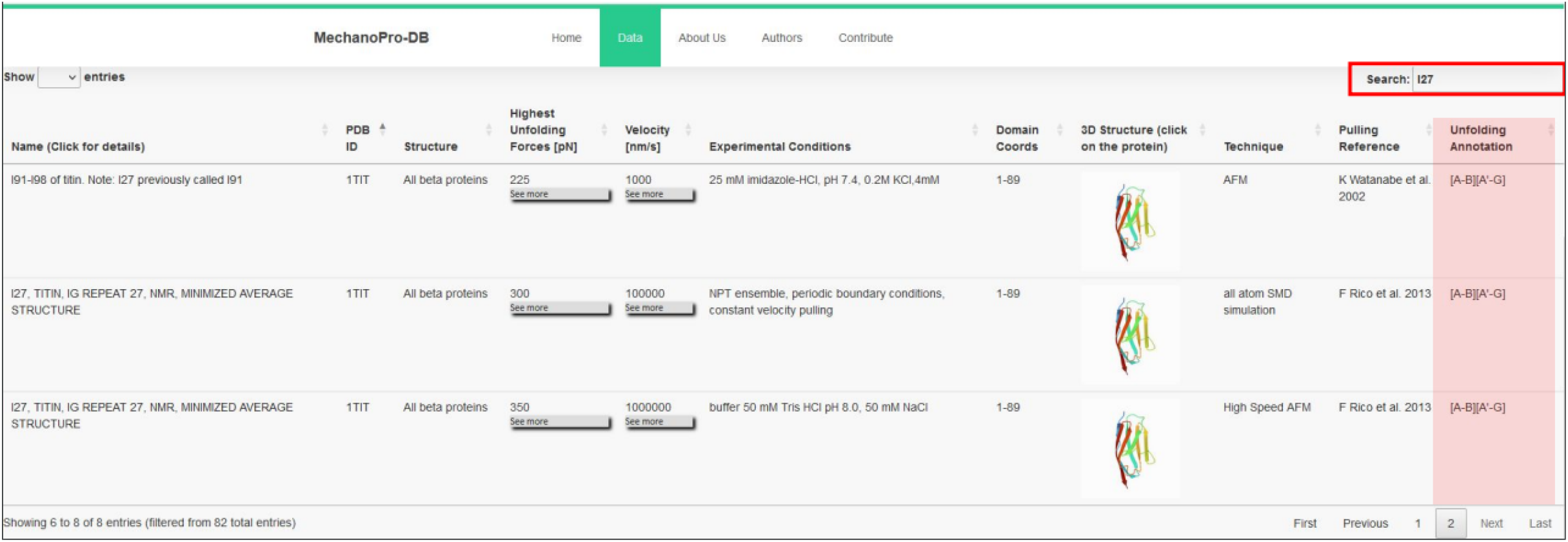
Screenshot showing the unfolding annotation of I27 module of titin after user search (red box). The annotation has been reported in the references displayed and is accessible within the ‘pulling reference’ column.

Each protein entry within MechanoProDB has a link to a dedicated page (**Figure 4**) which provides a curated visualization that enriches the understanding of the protein unfolding process. The main visual image within this page is a force-distance curve which shows examples of unfolding force measurements and structural transitions or, when available, the dynamic force spectrum, extracted from the original reference. Fits of theoretical models^25^ to the dynamic force spectra from the original publication enable us to include energy landscape parameters such as the free energy of a system (ΔG in kBT), the distance to the transition state along the reaction coordinate (here molecular extension, xβ in nm) and the rate at which the molecule transitions from one state to another (folded to unfolded) at zero applied force (*k*_*off*_ in 1/s).

**Figure 4:**
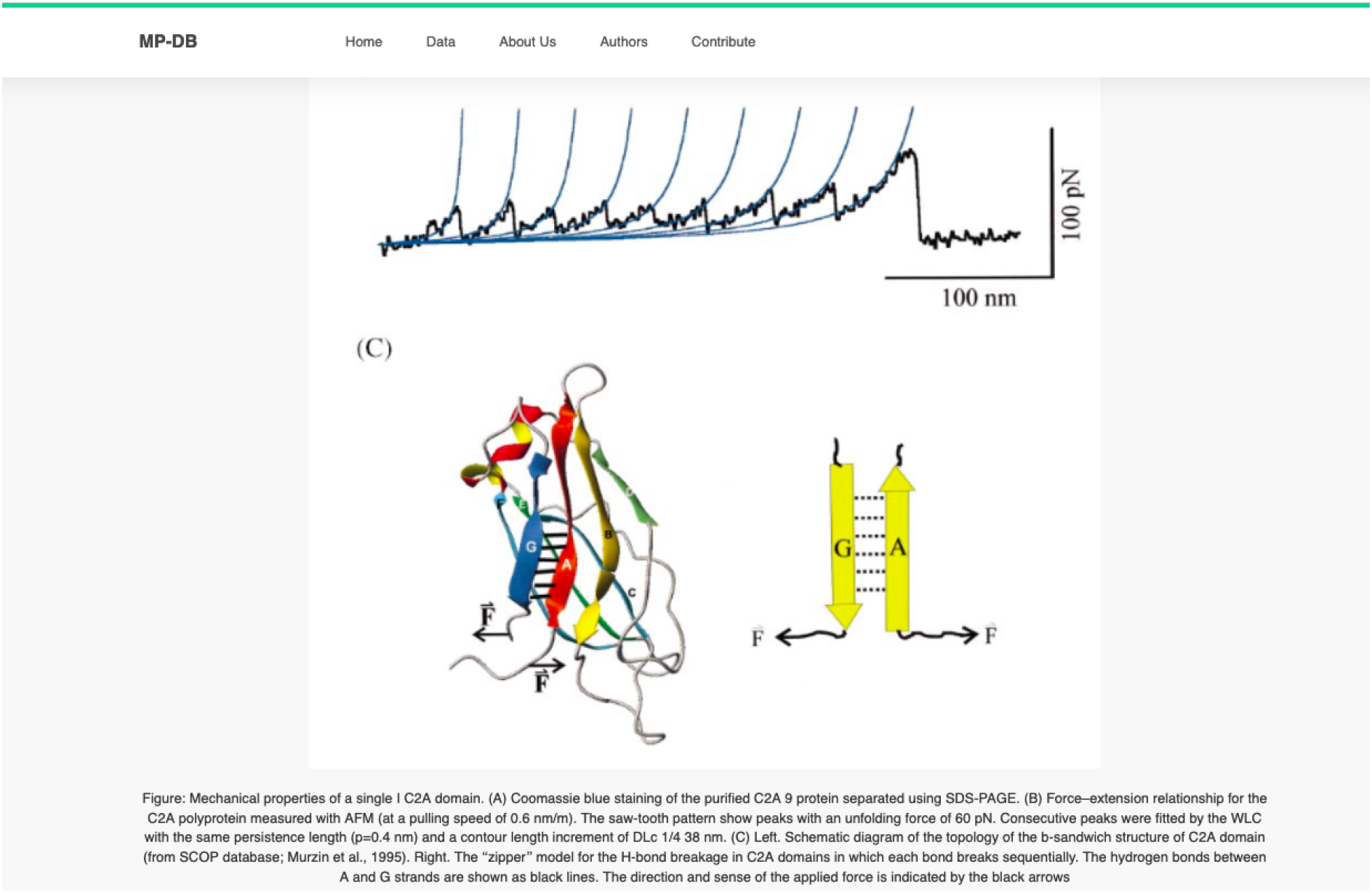
Snapshot of the detail page showing the synaptotagmin (C2A) domain. The figure, extracted from the original reference, displays the force-distance and the structural motif bearing the observed force^24^.

### Structure information and visualization

In addition to the mechanical properties, MechanoProDB provides structural information such as SCOP annotations which represent a systematic framework of assessing protein folds. According to this classification, proteins can belong to one of the following classes: all alpha proteins, all beta proteins, α+β protein, α/β protein and small proteins. This structural categorization is useful in illuminating the broader structural context of the proteins under investigation. SCOP annotations are derived from established structural methodologies, furnishing an insight into the fold architecture of proteins. This classificatory hierarchy can help in discerning shared structural motifs and evolutionary relationships among diverse protein entities. By incorporating SCOP annotations, MechanoProDB associates mechanical properties with structural information which facilitates a multidimensional understanding of protein mechanics.

Complementing these structural insights, we integrate other information including functional classification, the protein expression system and the protein organism. This information is extracted from the PDB. Functional classifications elucidates the roles and biochemical activities associated with the protein (e.g.: calcium binding, cell adhesion, transferase). The expression system information describes the conditions in which proteins were synthesized. The organismal information reports the source organism of the proteins, contextualizing their physiological relevance. By integrating diverse information derived from external resources such as SCOP and PDB, MechanoProDB reinforces its utility as a comprehensive resource. In MechanoProDB, each individual protein entry can be interactively visualized at sequence and structure level on a dedicated webpage, which can be reached by clicking on the structural representation of the protein on the main page (**Figure 5**). To achieve this, we have integrated the popular and versatile tool Mol*^19^ plugin, which offers an integrated perspective of protein sequence and its corresponding three dimensional structure. Notably, this plugin enables users to dynamically switch between diverse structural representations, from the familiar ribbon or cartoon representations to space filling models. On top of that, the plugin grants users the capability to elucidate solvent accessibility reinforcing the relevance of protein surface characteristics with respect to mechanical stability.

**Figure 5:**
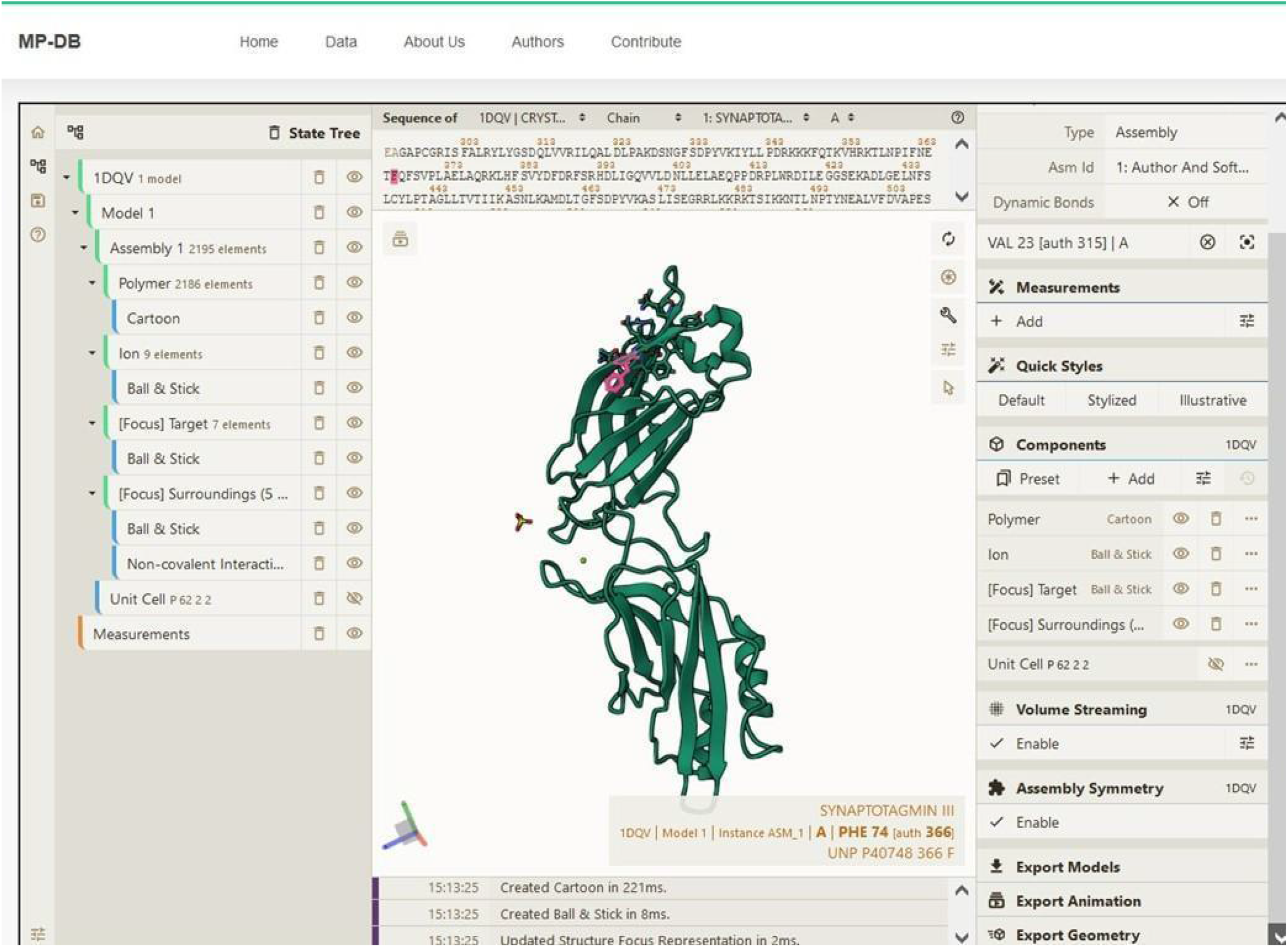
Snapshot showing the sequence and structure for the first domain of synaptotagmin (C2A) using the Mol^*^ plugin. This visualization enables users to examine sequence and three-dimensional structural conformation. In the example shown, Phenylalanine (F) is highlighted in both sequence and structure.

### Descriptive statistics of database content

In MechanoProDB, we incorporate interactive plots of relevant mechanical parameters stored in the database. On the main page, the first plot, shown in **Figure 6A**, represents unfolding forces versus pulling velocities of all entries. In addition to the force-velocity plot, a visualization of the percentage of each protein structural class, according to the SCOP annotation, is provided in a pie chart (**Figure 6B**). This percentage-based plot offers an immediate insight into the distribution of structural folds within the database. The third visualization (**Figure 6C)** shows boxplots that represent the distribution of unfolding forces across different protein structural folds as classified by SCOP. We can notice here that all beta proteins unfold at higher forces than all alpha proteins. Accordingly, the median unfolding force of proteins composed of β-sheets is ∼200 pN whereas that of α-helical proteins is ∼70 pN.

**Figure 6:**
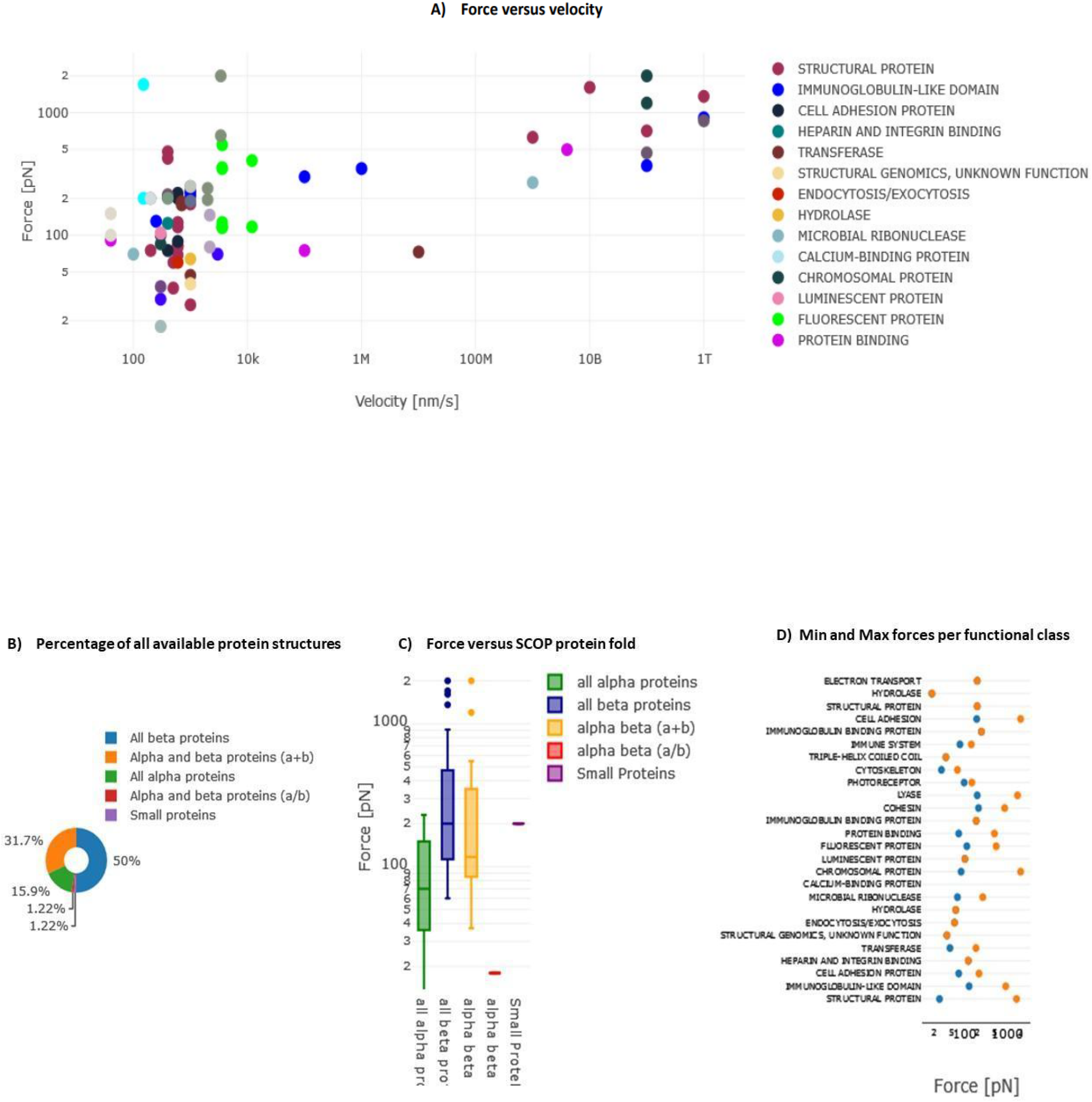
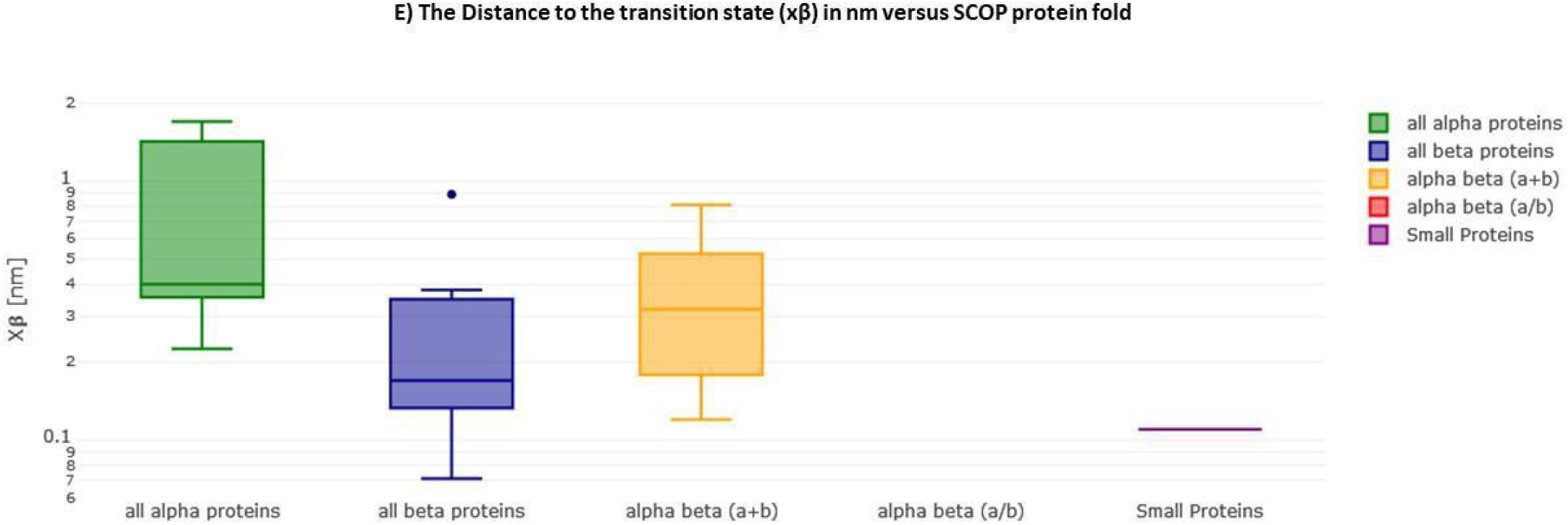
Interactive Plots in MechanoProDB. Illustration of the interactive plots within MechanoProDB: (A) Force versus velocity plot: Each data point represents a protein entry, colored by functional classification. Double-clicking a functional class isolates it, while a single click hides it. (B) Pie chart displaying the percentage of protein fold categories according to SCOP classification. (C) Boxplot depicting unfolding forces relative to protein fold, enabling cross-category comparisons. (D) Display of minimum and maximum forces categorized by functional class. Users can modify data representation by employing the pen tool, activated by hovering, see the red square on plot (A). (E) Boxplot showing the distance to the transition state with respect to SCOP structural classification.

A scatter plot (**Figure 6D**) illustrates the distribution of minimum and maximum unfolding forces within distinct functional classes. This elucidates the variance in mechanical forces across functional categories. The juxtaposition of force extremities within functional classes provides a representation of mechanical diversity within each class. Since unfolding forces depend on the applied pulling rate, energy landscape parameters provide more comparable information. **Figure 6E** displays boxplots of the distance to the transition state (xβ) with respect to SCOP structural classification, showing that, on average, the distance to the transition state of all beta proteins is shorter than that of all alpha proteins. The transition state corresponds to the highest energy point along the forced unfolding reaction coordinate. Boxplots representing distances to the transition state (xβ) versus SCOP classification show that all beta proteins cross the energy barrier at shorter distances (∼0.17 nm) than all alpha proteins (∼0.4 nm).

In MechanoProDB, all visual representations are interactive which empowers users to dynamically manipulate data displays. Users can isolate specific entities within the force-velocity plot, delve into force distributions across functional classes, explore mechanical variability based on SCOP folds and visualize prevalence of different protein structural classes. Users have also the possibility to change data representation by clicking on the pen that appears when hovering on each plot. In addition to the mentioned above plots.

### User contributions

In the spirit of encouraging collaborative engagement and expanding the utility of MechanoProDB, we enable user contributions and feedback (**Supp. Figure 2**). This allows users to actively participate in augmenting the database’s comprehensiveness.

Users desiring to insert new proteins with a solved 3D structure only need to provide the PDB-identifier as input. The PDB ID is supplied to an automated Python script that extracts complementary information. In scenarios where a given protein lacks a solved structure, users have to provide this information manually. Up to four unfolding forces can be submitted, each together with its corresponding pulling speed and/or loading rate. In case users possess a dataset surpassing four entries, MechanoProDB permits to submit data in various formats ranging from Comma-Separated Values (CSV) files to other tabulated formats (eg: dat, xls, txt,doc). A template xls file is freely-available for download from the user contribution page. Finally, MechanoProDB provides written tutorials and videos are available to familiarize users with the different services provided by the database.

## DISCUSSION & CONCLUSIONS

The principal aim of MechanoProDB is to provide a comprehensive compilation of mechanical properties of proteins. This compilation is meticulously extracted from published articles. The information stored within the database includes a variety of experimental and computational techniques. Experimental data includes atomic force microscopy, optical and magnetic tweezers data obtained in the force spectroscopy mode. Complementary to this, our database integrates computational data derived from steered molecular dynamics simulations data both from all-atom and coarse-grained modes.

The development of MechanoProDB represents a significant contribution to the growing field of molecular mechanobiology and single molecule biophysics. It addresses a critical need for a centralized and freely accessible resource that stores and makes available the mechanical properties of proteins, otherwise dispersed and difficult to localize by non-experts. MechanoProDB provides an expansive dataset of mechanical properties with high-quality manual annotations of the unfolding pathways. The inclusion of SCOP annotations and functional classifications enhances and advances our understanding of the relationship between protein sequence, structure and mechanics which contextualizes these mechanical profiles within the broader structural landscape.

The interactive visualization offered by MechanoProDB empowers users to analyze unfolding forces with experimental conditions, functional classes, and protein folds. The boxplots showing the unfolding forces with respect to SCOP annotations consolidate prior findings^6^ regarding the correlation between unfolding forces of proteins and structural classes (all beta proteins versus all alpha proteins). Indeed, proteins composed of β-sheets unfold at higher forces than those composed of α-helixes, a previously suggested generalization now quantitatively confirmed. The integration of the molecular visualization package Mol* further improves analytical capabilities of sequence and structure, adding an extra layer of insight by offering a simultaneous sequence and structural visualization, which may help understanding structural mechanics.

A notable advantage of MechanoProDB is its provision of detailed and accurate information about the mechanical unfolding of proteins from existing published works using experimental and computational approaches. Compared to the only other existing database, BSDB^13^, MechanoProDB collects a more complete scope of information, including mechanics, structure and function of experimental and computational data from the literature. Additionally, compared to BSDB, MechanoProDB goes beyond unfolding forces, velocities, and CATH references by providing a broader range of data and comprehensive unfolding annotations, including unfolding pathways, experimental conditions, and energy landscape parameters such as free energy, distance to transition state and intrinsic unfolding rate. Moreover, MechanoProDB is a dynamic platform that allows users to plot and visualize the stored data and gives them the possibility to contribute their own data and feedback. These functionalities facilitate rigorous and insightful research, positioning our database as an important, community-shared platform for scientists interested in protein mechanics.

We hope that MechanoProDB will enable a deeper exploration of the relationships between sequence, structure, function and mechanics. By stimulating user contributions and feedback, MechanoProDB provides a collaborative platform in which researchers collectively contribute to advancing the understanding of protein (un)folding mechanics. This participative model emphasizes the fundamental principles of open science, enabling dynamic exchange of knowledge and insights within the protein research community. By fostering collaboration and elevating comprehensiveness of protein mechanical data, MechanoProDB may emerge as a useful resource in the field of experimental and computational biomechanics. We believe that this resource will make a significant contribution to the understanding of protein mechanics and inspire further investigations into the complex field of mechanobiology.

## DATA AVAILABILITY

MechanoProDB is freely available at https://mechanoprodb.ibdm.univ-amu.fr. To ensure optimal data utilization, MechanoProDB, offers a dedicated tutorial page within the website. This page serves as a guide, elucidating the functionalities and features of the database. Thereby, MechanoPrDB empowers users to effectively navigate and explore its potential. Within the tutorial page, we also provide an informative video tutorial. This visual resource illustrates the database’s interface, features and interactions. All code is freely accessible on GitLab via the following link: https://gitlab.com/fm4b_lab/mechanoprotein-db

## ACKNOWLEDGEMENTS

We acknowledge Dr. Rafael Tapia-Rojo for contributing with data.

We thank Dr. Mar Benavides and Dr. Jean-Luc Pellequer for inspiring comments.

