## supplementary file for "MechanoProDB: A Web Based Database for Exploring the Mechanical Properties of Proteins"

**SUPPLEMENTARY DATA**

- Tutorial video.
- **Table S1**: percentage of proteins unfolding at 1,2,3,4 and 5 steps:

| | all alpha proteins | all beta proteins | α + β proteins | α / β proteins | small proteins |
|---|---|---|---|---|---|
| $N - U$ | 42.8% | 82.5% | 58% | 0 | 100% |
| $N - I - U$ | 14.2% | 12.5% | 33% | 0 | 0 |
| $N - I - I - U$ | 14.2% | 0 | 0 | 100% | 0 |
| $N - I - I - I - U$ | 28.5% | 5% | 0 | 0 | 0 |
| $N - I - I - I - I - U$ | 0 | 0 | 8% | 0 | 0 |

Table S1 legend: 42.8% of all alpha proteins unfold in 1 step. all beta proteins are more likely to unfold in one step.

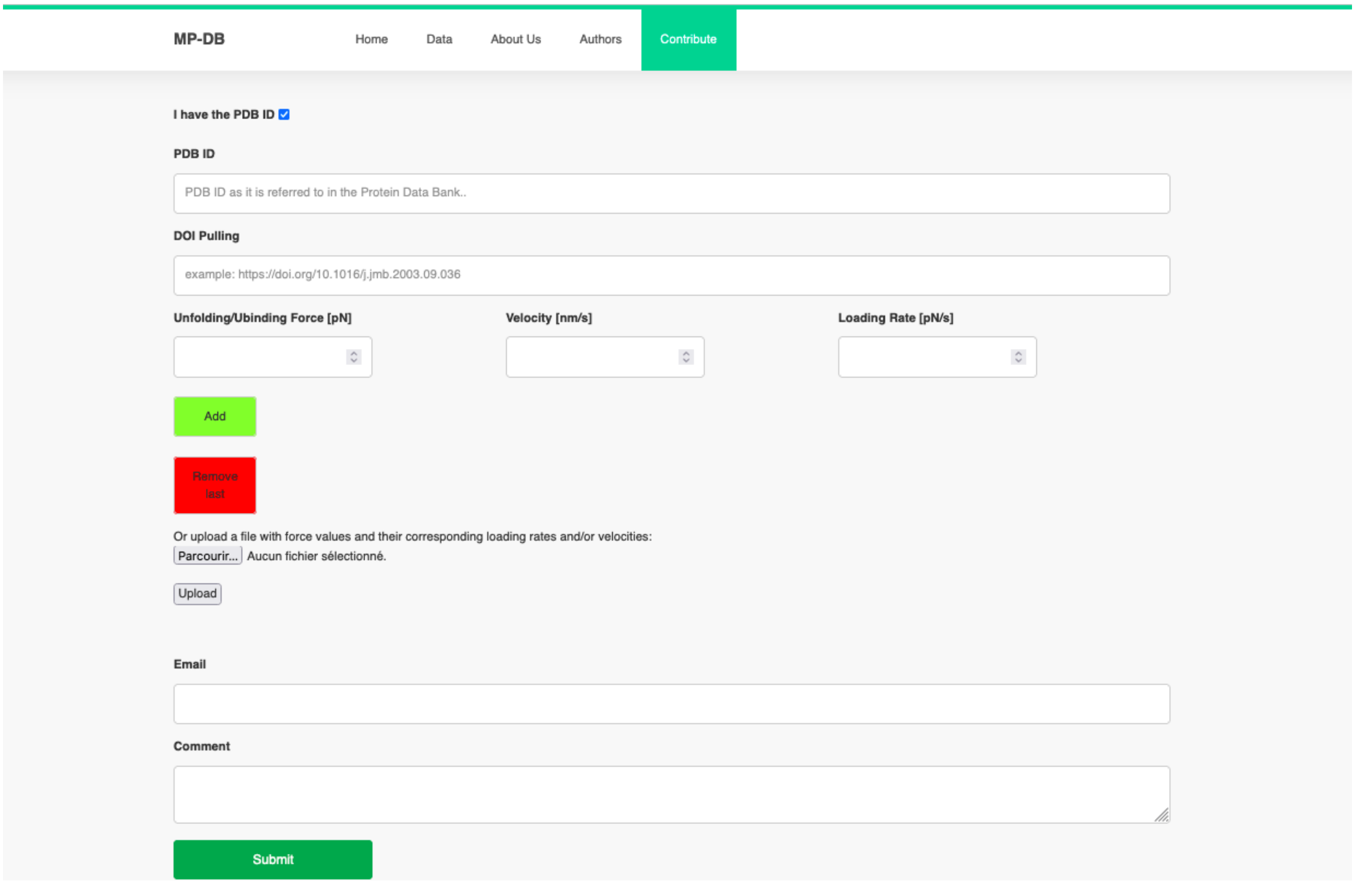


**Supp. Figure 2**: Illustration of MechanoProDB's upload page for contributing to MechanoProDB. Users can actively participate by submitting protein entries with crystal structures, providing PDB IDs for automated data extraction. For proteins without PDB accession code, comprehensive datasets can be contributed.
